# Characterisation of protective vaccine antigens from the thiol-containing components of excretory/secretory material of *Ostertagia ostertagi*

**DOI:** 10.1101/2023.12.08.570749

**Authors:** Daniel R. G. Price, Philip Steele, David Frew, Kevin McLean, Dorota Androscuk, Peter Geldhof, Jimmy Borloo, Javier Palarea Albaladejo, Alasdair J. Nisbet, Tom N. McNeilly

## Abstract

Previous vaccination trials have demonstrated that thiol proteins affinity purified from *Ostertagia ostertagi* excretory-secretory products (*O. ostertagi* ES-thiol) are protective against homologous challenge. Here we have shown that protection induced by this vaccine was consistent across four independent vaccine-challenge experiments. Protection is associated with reduced cumulative faecal egg counts across the duration of the trials, relative to control animals. To better understand the diversity of antigens in *O. ostertagi* ES-thiol we used high-resolution shotgun proteomics to identify 490 unique proteins in the vaccine preparation. The most numerous ES-thiol proteins, with 91 proteins identified, belong to the sperm-coating protein/Tpx/antigen 5/pathogenesis-related protein 1 (SCP/TAPS) family. This family includes previously identified *O. ostertagi* vaccine antigens *O. ostertagi* ASP-1 and ASP-2. The ES-thiol fraction also has numerous proteinases, representing three distinct classes, including: metallo-; aspartyl– and cysteine proteinases. In terms of number of family members, the M12 astacin-like metalloproteinases, with 33 proteins, are the most abundant family in *O. ostertagi* ES-thiol. The *O. ostertagi* ES-thiol proteome provides a comprehensive database of proteins present in this vaccine preparation and will guide future vaccine antigen discovery projects.

## 1. Introduction

The gastrointestinal nematode *Ostertagia ostertagi* causes serious health and welfare issues as well as substantial economic losses in cattle. Control of this parasite is reliant on the use of anthelmintics, but the widespread and prolonged use of these drugs has resulted in the selection of resistant worms (Bartley et al., 2021; Cotter et al., 2015; Edmonds et al., 2010). Alternative control approaches, such as vaccination, are needed to provide robust and sustainable control of these parasites. Previous *O. ostertagi* vaccination trials have included immunisation of cattle with live attenuated larvae (Bürger and Pfeiffer, 1969), with somatic antigens (Herlich and Douvres, 1979) or unfractionated ES products (Hilderson et al., 1995). None of these approaches have resulted in significant levels of protection against *O. ostertagi* infection. However, vaccination of calves with thiol-binding proteins that have been affinity purified from adult *O. ostertagi* ES (*O. ostertagi* ES-thiol) results in statistically significant levels of protection, with a 56 – 60 % reduction in mean cumulative faecal egg counts in vaccinates, relative to control, unvaccinated cattle (Geldhof et al., 2004, 2002).

The *O. ostertagi* ES-thiol fraction contains *Ancylostoma* secreted proteins (ASPs) and, of these, Oo-ASP-1 and Oo-ASP-2 have been identified as protective antigens in their native form (Geldhof et al., 2002, 2003a; Meyvis et al., 2007a). Other ASPs from parasitic nematodes have also shown promise as vaccine antigens; for example, ASPs from the hookworms *Ancylostoma caninum* (Ghosh et al., 1996; Ghosh and Hotez, 1999; Hilderson et al., 1995) *and A. ceylanicum* (Goud et al., 2004; Mendez et al., 2005) provide significant levels of protection against parasite challenge and the recombinant cocktail vaccine against *Teladorsagia circumcincta* also contains an ASP (Nisbet et al., 2013). The ASPs belong to an evolutionary conserved group of secreted proteins from the SCP/TAPS (sperm-coating protein/Tpx/antigen 5/pathogenesis-related protein 1) family, and are present in diverse eukaryotic organisms, including parasitic nematodes. The SCP/TAPS are secretory proteins, characterised by the presence of either one or two cysteine-rich, antigen 5, and pathogenesis-related 1 (CAP) domains (CAP, InterPro domain IPR014044). The first reported SCP/TAPS protein was Ac-ASP-1 from the secretome of the hookworm, *A. caninum* (Hawdon et al., 1996). Since then, SCP/TAPS proteins have been identified in the secretome of many parasitic nematodes, often as part of large expanded families, and are likely to play an important role in parasitism (International Helminth Genomes Consortium, 2019; Viney, 2017).

ASPs from the adult *O. ostertagi* ES-thiol fraction have shown great promise for vaccine development (Geldhof et al., 2008, 2003b, 2002; González-Hernández et al., 2016; Meyvis et al., 2007b). Purified native *O. ostertagi* ASP-1 protected cattle against homologous challenge, with greater levels of protection compared to animals vaccinated with the ES-thiol fraction (Meyvis et al., 2007b). However, the development of a recombinant version of Oo-ASP-1 that protects against *O. ostertagi* infection has been a challenge. Initial experiments demonstrated that recombinant Oo-ASP-1 produced in yeast failed to protect animals against homologous challenge, even though the protein antigen was correctly folded and had the same three-dimensional protein shape as the native molecule (Borloo et al., 2013; González-Hernández et al., 2016). This failure in protection is likely due to the recombinant expression system being unable to reconstruct native structures of the worm protein, including post-translational modifications, such as glycosylation. Recent success has been achieved be production of recombinant oO-ASP-1 produced in plants, were the recombinant protein is decorated with worm-like glycans present on the native molecule. This refined recombinant protected vaccinated calves from *O. ostertagi* infection (personal communication, Geldhof *et al*. 2023). Thus, for a successful recombinant antigen, both the selection of antigen, as well as ability to produce a recombinant version that reflects the native molecule is critical for development of a successful vaccine.

In addition to ASPs, it is clear that the *O. ostertagi* ES-thiol protein fraction contains other protective antigens (Meyvis et al., 2007a) and these may offer a source of additional vaccine antigens. This is demonstrated by a study in which *O. ostertagi* ES-thiol was fractionated into three sub-fractions, with each protein fraction providing significant levels of protection against *O. ostertagi* infection, relative to control animals (Meyvis et al., 2007a). Although the ES-thiol fraction is a potential source of other protective vaccine antigens, the protein diversity within this fraction is largely unknown. Therefore, the objective of this present study was to apply high-resolution shotgun proteomics to discover the diversity of proteins within *O. ostertagi* ES-thiol.

## 2. Materials and Methods

### 2.1 Ethics Statement

All animal procedures were performed at Moredun Research Institute (MRI), UK under project licenses 70/7914 and P23F688B4 as required by the UK Animals (Scientific Procedures) Act 1986, with ethical approval from MRI Animal Welfare and Ethical Review Body. Animals were maintained at MRI under conditions designed to exclude accidental infection with helminth parasites and were confirmed helminth-naïve at the start of the experiments.

### 2.2 Production of parasite material

To generate *O. ostertagi* adult worms for excretory-secretory product (ESP) generation, helminth-free Holstein-Friesian male calves (4-10 months old) were challenged with 50,000 infective third-stage (L3) *O. ostertagi* larvae (isolate MOo2). At 21 days post infection (dpi) animals were euthanised and *O. ostertagi* adults were recovered from the gastric stomach (abomasum) of individual animals. In brief, the abomasum was opened along the greater curvature and washed in physiological saline (0.9 % NaCl (w/v)). The abomasal wash sample was poured over a 212 µm sieve and floated on physiological saline at 37 °C for four hours and *O. ostertagi* parasites that migrated through the sieve were collected. Following washing, the abomasum was pinned to a polystyrene board and floated on physiological saline at 37 °C for four hours and *O. ostertagi* parasites that migrated out of the tissue were collected.

### 2.3 Preparation of *O. ostertagi* Excretory-Secretory Products

At 21 dpi, *O. ostertagi* adults were recovered from the abomasum of infected calves, as described above. For collection of ESPs, harvested parasites were washed three times in phosphate-buffered saline (PBS) before culturing at 37 °C with 5% CO_2_ in RPMI 1640 medium (Invitrogen, Carlsbad, CA, USA) containing: 20 mM HEPES pH 7.5; 10 mM L-glutamine; 1000 U/ml penicillin; 1000 μg/ml streptomycin; 200 μg/ml gentamycin and 5 μg/ml amphotericin B. Culture supernatants were harvested every 24 h and replaced with fresh media up to 96 h. At each time point the viability of the parasites was confirmed on the basis of structural integrity and motility. The culture supernatants containing *O. ostertagi* ESPs were filtered using Millipore Express 0.22 µm PES 500ml filters and stored at –70 °C. Prior to chromatography, ESPs were concentrated, and buffer exchanged into PBS using Amicon Ultra-15 columns (Millipore) with a 10-kDa molecular mass cut-off. Protein concentrations were determined by BCA assay (Thermo Fisher Scientific) with BSA standards.

### 2.4 *O. ostertagi* Excretory-Secretory Product fractionation

Thiol-binding proteins were purified from *O. ostertagi* ESPs according to the method of Geldhof *et al*., 2002 (Geldhof et al., 2002). In brief, a 5 ml thiol-sepharose column (Cytiva) was equilibrated in 5 column volumes of running buffer (10mM Tris, 0.5M NaCl, pH7.4). Immediately prior to chromatography, approximately 5 mg of *O. ostertagi* day 21 ESPs was reduced by incubation with 2.5 mM DTT (final concentration) for 30 min at 37°C and excess DTT was removed by desalting on a HiPrep 26/10 column. The reduced desalted ESP sample was applied to the thiol-sepharose column at 0.1 ml/min, the column was then washed with 10 column volumes of running buffer. Bound proteins were eluted with running buffer plus 50 mM DTT. Equilibration, washing and elution steps carried out at 0.8 ml/min. Eluted fractions were desalted on a HiPrep 26/10 column using 10 mM Tris-HCl pH 7.4 as the running buffer. ES-thiol proteins were then concentrated using Amicon Ultra-4, 10kDa centrifugal concentrators. Protein concentration was determined by BCA protein assay (Thermo Fisher Scientific) with BSA standards and proteins analysed by reducing SDS-PAGE gel.

### 2.5 SDS-PAGE

Proteins were analysed by SDS-PAGE using the NuPAGE^®^ electrophoresis system (Thermo Fisher Scientific, Waltham, MA, USA). Briefly, protein samples were prepared in NuPAGE LDS sample buffer with reducing agent and heated to 70°C for 10 min prior to loading on NuPAGE 4–12% Bis-Tris gels. Gels were run in MES SDS running buffer and stained with SimplyBlue™ SafeStain (Thermo Fisher Scientific, Waltham, MA, USA).

### 2.6 Vaccination Trials

A series of four vaccine trials were performed between 2017 and 2021 (Trials 1-4) to test the efficacy of an ES-Thiol-based vaccine against *O. ostertagi* challenge. The design of each trial is summarised in Supplementary Table 1. For Trials 1 and 2, calves were immunised with 30 µg ES-Thiol plus Quil A^®^ (Brenntag Biosector) adjuvant or Quil A^®^ only (adjuvant control) on days 0 (V1), 21 (V2) and 42 (V3) via the intramuscular route. For Trials 3 and 4, calves were immunised with 15 µg ES-Thiol plus Vax Saponin^®^ (Guinness) adjuvant or Vax Saponin^®^ only (adjuvant control) on days 0 (V1), 21 (V2) and 42 (V3) via the subcutaneous route (Trials 3-4). Calves were between 4 to 7.5 months old at first vaccination and were randomly allocated to either vaccine or adjuvant control groups balancing for age, sex, breed, and source farm. At V3, calves were either trickle infected with 25,000 *O. ostertagi* L3 isolate MOo2 (1,000 L3 per day *per os*, 5 days a week for 5 weeks) (Trials 1-2) or challenged once with an oral bolus of 50,000 *O. ostertagi* L3 isolate MOo2 (Trials 3-4). Blood samples were collected via jugular venepuncture at days 0 and 28 for Trial 1-2 and days 0 and 42 for Trials 3-4 to determine serum levels of ES-Thiol specific antibodies as detailed in Section 2.7. Faecal samples were collected twice weekly post-L3 challenge for *O. ostertagi* faecal egg count measurements. Calves were euthanized on day 91 relative to V1 for Trial 1, day 98 for Trial 2 and day 76 for Trials 3 and 4, for worm burden estimation as detailed in Section 2.7.

### 2.7 Parasitological Measurements

Faecal egg count (FEC) analysis was performed twice weekly on up to 10 g of faeces collected from each animal from day 10 post-challenge as previously described (Christie and Jackson, 1982). FEC over time was analysed by fitting a Poisson generalised additive mixed model (GAMM) with FEC as response, group as fixed effect and animal nested within trial as random effect. Separate smoothing splines were considered for each group in order to describe their temporal pattern while allowing for heterogeneous variances and autocorrelation. Cumulative FEC (cFEC) was estimated over the course of each trial by calculating the area under the FEC curves for each animal using the composite trapezoid rule (Taylor et al., 1997). Differences in cFEC between vaccinated and control groups was analysed using a negative binomial generalised linear mixed model (NB GLMM), including cumulative FEC as response variable, treatment group as fixed effect, and trial as random effect. Enumerations of abomasal worm burdens were carried out at post-mortem following standard techniques (Halliday et al., 2007; Jackson et al., 1984). Differences in total worms (luminal and mucosal) between vaccinates and controls was analysed using negative binomial GLMs accounting for data over-dispersion by estimating a dispersion parameter.

### 2.8 Measurement of vaccine-specific antibody response

Serum IgG1 and IgG2 levels against *O. ostertagi* ES-thiol were determined by ELISA on the day of the first vaccination (V1, day 0) and V2 +7days (Trials 1 & 2) and on the day of the first vaccination (V1, day 0) and V3 (day 42) for Trials 3 & 4. Ninety-six-well plates were coated with 50 µl/well of ES-thiol diluted in 0.05 M carbonate buffer (pH 9·6) at a concentration of 1 µg/ml. Non-specific binding was blocked with 3 % (w/v) fish gelatine in PBS (Sigma G7765). Individual cattle serum samples were applied in duplicate at 1:1000 (IgG1) or 1:500 (IgG2). Secondary antibodies used were BioRad MCA627GA mouse anti-bovine IgG1 (1:1000) and BioRad MCA626GA mouse anti-bovine IgG2 (1:2000) followed by the tertiary conjugate rabbit anti-mouse Ig conjugated to horseradish peroxidase, Dako P0260, (1:1000). Immunoreactivity was visualised by incubation with OPD substrate (SigmaFast OPD, P9187) at room temperature for 5-10 minutes, before stopping the reaction by addition of 25 µl 2.5 M H_2_SO_4_. All dilutions were made in PBS, 0.5 % Tween80, 0.5 M NaCl and incubation steps carried out for 1 hr at 37°C. Plates were washed six times between each step in PBS, 0.05 % Tween 20 using a Bio-Tek ELx405 plate washer. A positive control sample was included in each plate to account for inter-plate variation. Optical density was measured at 492 nm using a Molecular Devices SpectraMax ABS Plus plate reader. Statistical analysis was performed in GraphPad Prism by using two-way ANOVA with Tukey’s multiple comparison test.

### 2.9 PacBio Iso-Seq library preparation and sequencing

At 21 dpi, *O. ostertagi* adults were recovered from the abomasum of infected calves, as described above. Male and female *O. ostertagi* parasites were viewed using a dissecting microscope and picked using a fine needle into separate tubes. Parasites were snap-frozen in liquid nitrogen and stored at –70 °C until RNA isolation. Total RNA was isolated from *O. ostertagi* male and female worms separately using RNeasy mini kits (Qiagen) with a DNaseI digest, according to the manufacturer’s instructions. Total RNA was quantified using a NanoDrop^TM^ One spectrophotometer and quality assessed with an Agilent 2100 Bioanalyser using an RNA chip (Agilent Technologies). For library construction all RNA samples had a RIN value >7. Library preparation and sequencing were conducted at the University of Liverpool, Centre for Genomic Research. The sample was initially treated with Teloprime full length cDNA amplification kit v2 (Lexogen GmbH) following the manufacturer’s protocol. This involved first strand cDNA synthesis by denaturation of sample (1000 ng) at 70°C with primer RTP and after cooling to 37°C mixing with reverse transcriptase and incubating at 46°C for 50 minutes. The sample was cleaned using the supplied columns. The sample was then ligated to a cap-dependent linker for 3 hours at 25°C and cleaned again with the same columns as before. The sample was incubated with 2^nd^ strand synthesis mix for 90 sec at 98 °C, 60 sec at 62 °C, and 72 °C for 5 minutes with a hold at 10 °C. The sample was column cleaned again. Half of the sample was amplified thereby making ds-cDNA for 15 cycles (1× (95.8 °C 30 sec, 50 °C 45 sec, 72 °C 20 minutes), 14× (95.8 °C 30 sec, 62 °C 30 sec, 72 °C 20 minutes) and cleaned with a column as before. The amplified cDNA products were made into SMRTbell template libraries according to the Iso-Seq protocol by Pacific Biosciences. Sequencing was performed on the PacBio Sequel System, and 1 SMRT Cell was run for each sample with a movie run-time of 600 min for each SMRT Cell.

### 2.10 Sequence Processing

For each sample, the CCS module of the IsoSeq v3 program (https://github.com/PacificBiosciences/IsoSeq) was used, with default parameter settings, to generate circular consensus sequence (CCS) reads from the sub-reads generated from each sequencing run. The CCS module provides full-length reads spanning entire transcript isoforms all the way from the polyA-tail to the 5′ end. High quality isoforms were further clustered using CD-HIT version 4.6 (Fu et al., 2012) with the following parameters: –c = 0.97 –G = 0 –aL = 0.8 –AL = 100 –aS = 0.99 –AS = 30 –g = 1. Open reading frames (ORFs) were predicted using ANGEL ORF prediction software (https://github.com/PacificBiosciences/ANGEL) and annotated using Blast2GO through OmicsBox software (ver. 2.1, BioBam Bioinformatics Solutions).

### 2.11 Transcriptome Completeness

Transcriptome completeness was determined using Benchmarking Universal Single-Copy Orthologs (BUSCO; v5.2.2; –mode proteins –lineage metazoa_odb10) gene completeness metrics (Simão et al., 2015).

### 2.12 Proteomic analysis of *O. ostertagi* ES-thiol

#### 2.12.1 Liquid digest

ES-thiol samples were digested using an S-Trap micro column (Protifi) kit, following the manufacturer’s instructions. Briefly, 1-2 µg of *O. ostertagi* ES-thiol was mixed with the supplied 2X SDS lysis buffer, proteins were reduced with 5 mM Tris (2-carboxyethyl) phosphine (TCEP) and alkylated with 20 mM methyl methanethiosulfonate (MMTS). Proteins were then acidified with 2.5% phosphoric acid and bound to the S-Trap column in 6 volumes of S-Trap binding/washing buffer (100mM TEAB in 90% Methanol). The column was centrifuged (4000 x*g* for 30 s) and then washed a further 4 times in wash buffer. Proteins were digested using sequencing grade trypsin (in 50 mM TEAB) in a 1:10 ratio to trapped protein for 16 hrs at 37°C. Tryptic peptides were eluted in three steps, firstly, 40 μl 50 mM triethylammonium bicarbonate (TEAB) buffer followed by 40 μl 0.2 % formic acid (FA), and finally 40 μl 50 % acetonitrile (ACN) in water and 0.2 % FA. Elutions were pooled and dried in a vacuum drier. Dried peptides were reconstituted in 20 µl of 0.1% FA prior to analysis by Mass Spectrometry.

#### 2.12.2 Gel piece digest

A single SDS-PAGE gel lane containing *O. ostertagi* ES-thiol proteins was excised and sliced horizontally from top to bottom to yield a series of 26 equal gel slices. The resulting gel slices were destained with 50mM ammonium bicarbonate in 50% acetonitrile (ACN). Proteins were reduced with 10mM DTT in 100mM ammonium bicarbonate and alkylated with 55mM Iodoacetamide in 100mM ammonium bicarbonate was added and placed in the dark for 30min. Proteins were digested with trypsin (Promega Porcine trypsin, 10 ng/µl trypsin in 25 mM ammonium bicarbonate) at 37°C overnight.

#### 2.12.3 LC-MS/MS analysis

Liquid chromatography−tandem mass spectrometry (LC−MS/MS) analysis was performed using an Ultimate 3000 nano-LC system coupled to a Q-Exactive Plus mass spectrometer (Thermo Fisher Scientific). The LC system was equipped with an Acclaim Pepmap nano-trap column (C18 PepMap100, 300 μm × 5 mm, 5 μm, 100 Å, Thermo Fisher Scientific) and an Acclaim Pepmap RSLC analytical column (C18, 100 Å, 75 μm×50 cm EasySpray, Thermo Fisher Scientific). The tryptic peptides were loaded onto the trap column at an isocratic flow of 5 μL/min of 0.05% v/v trifluoroacetic acid for 6 min, peptides were then separated on the Easy Spray analytical column. The eluents were 0.1% v/v formic acid (solvent A) in H_2_O and 80% v/v CH_3_CN in 0.1% v/v formic acid (solvent B). A 65 min gradient was used at 300 nl/ min from (i) 0–5 min, 5% B; (ii) 5-40 min, 5–40% B; (iii) 40–45 min, 40–80% B; (iv) 45-50 min, 80% B; (v) 50-65 min, 80–5% B.

Data were acquired in positive mode using a data dependent approach, MS1 scans were acquired at 70,00 resolution over a mass range of m/z 380-1500 with an AGC of 3e6 and a maximum IT of 100ms. In each cycle, the 10 most intense ions with charge states of ≥2 and intensity thresholds of ≥2.0e5 were selected for MS/MS and subjected to high-energy collision dissociation (HCD) at a normalized collision energy of 30%. The isolation window was set at 1.2 m/z, at a resolution of 17,500, an AGC target of 1e5 and a maximum IT time of 100 ms. The dynamic exclusion time was set to 30 s.

#### 2.12.4 Proteomic Data Analysis

The MS raw data was processed using the Proteome Discoverer platform (version 2.4, Thermo Fisher Scientific) and Sequest HT algorithm. The MS data was searched against the translated *O. ostertagi* male and female Iso-Seq database. A maximum of two missed tryptic cleavages were allowed for and the minimum peptide length was six amino acids. The oxidation of Methionine and protein N-terminal acetylation were set as variable modifications, while carbamidomethylation of Cysteine was set as a fixed modification. MS and MS/MS ion tolerances were set at 10 ppm and 0.02Da, respectively. A maximum false discovery rate (FDR) of 1% at both the peptide and the protein levels was set and only proteins identified with ≥2 peptides were accepted. The lists of accepted proteins from the 26 Gel-LC slices were compiled into one list using Proteome Discoverer. The Gel-LC output was then merged with the output from 3 liquid digest loadings to give a final list of identified and accepted proteins.

#### 2.12.5 Protein database compilation

To generate a PacBio Iso-Seq derived database for MS searching, we grouped high-quality isoforms from *O. ostertagi* male (15,521 isoforms) and female (13,387 isoforms) and clustered isoforms using CD-HIT version 4.6 (Fu et al., 2012) with the following parameters: “-c = 1.0 –n = 5 –G = 1 –g = 1 –b = 20 –l = 10 –s = 0.0 –aL = 0.0 –aS = 0.0”. The resultant protein coding sequence database, containing 22,698 unique isoforms, formed the reference proteome.

#### 2.12.6 Annotation of protein sequences

For annotation, the FASTA protein sequences were imported into OmicsBox version 3.1 (BioBam) and annotated using blastp search against NCBI nr database (E-value ≤ 1.0 × 10^−3^). Subsequent GO mapping was performed using the Blast2GO mapping against the latest version of the GO database to obtain the functional labels (Götz et al., 2008). Prediction of transmembrane domains and signal peptide sequences was done using the TOPCONS prediction server (Tsirigos et al., 2015).

### 2.13 Phylogenetic analysis of CAP domain proteins

For phylogenetic analysis, ninety-one *O. ostertagi* CAP domain proteins identified in the *O. ostertagi* ES-thiol fraction were analysed in a framework of previously characterised CAP domain containing proteins from clade V parasitic nematodes. This included four ASPs from *O. ostertagi* (Oo-ASP-1 [CAD23183.1] from (Vercauteren et al., 2003); Oo-ASP-2 [CAD56659.1] from (Geldhof et al., 2003a); Oo-ASP-3 [CAO00416.1] and Oo-ASP-4 [CAO00417.1] from (Visser et al., 2008)) and one ASP from *T. circumcincta* (Tci-ASP-1 [CBJ15404.1] from (Nisbet et al., 2010)). In addition, we included nine ASPs from *Ancylostoma caninum*, which have become the canonical set of hookworm ASPs (Acan-NIF [AAA27789.1] from (Moyle et al., 1994); Acan-PI [AAK81732.1] from (Del Valle et al., 2003); Acan-ASP-1 [Q16937.1] from (Hawdon et al., 1996); Acan-ASP-2 [AAC35986.1] from (Hawdon et al., 1999); Acan-ASP-3 [AAO63575], Acan-ASP-4 [AAO63576], Acan-ASP-5 [AAO63577], Acan-ASP-6 [AAO63578] from (Zhan et al., 2003) and Acan-ASP-7 [AEJ86344] from (Datu et al., 2008)). Further CAP domain proteins were retrieved from the proteomes of *H. contortus* (Doyle et al., 2020) (bioproject: PRJEB506; 229 CAP domain proteins) and *T. circumcincta* (bioproject: PRJNA72569; 203 CAP domain proteins) based on the presence of Interpro CAP domain (IPR014044) using an E-value threshold of <10^-3^. The retrieved protein set contained a total of 537 CAP domain proteins, the full list is available in Supplementary File 1.

For sequence alignment we used a single domain CAP protein from *O. ostertagi* (transcript/11662) that had been trimmed to its CAP domain (amino acids 129 – 274) to extract an alignment of CAP domains from the protein set. Briefly, CAP domains were extracted using jackhmmer (from HMMER package 3.3.2) which was run for 10 iterations, or to convergence before then, at maximum sensitivity, and with E-value thresholds of 10^−6^, with the following parameters “-N 10 –E 1e-6 –-domE 1e-6 –-incE 1e-6 –-incdomE 1e-6 –max –A <output alignment file>”. The resulting protein domain sequences were extracted from the output alignment file and realigned with MUSCLE 5.1 (Edgar, 2021) and trimmed with ClipKit 1.3.0 (Steenwyk et al., 2020). Aligned and trimmed CAP domain sequences were used to create maximum-likelihood phylogenies using FastTree 2.1.11 (Price et al., 2010) using the LG amino acid substitution model and the reliability of the tree was tested using the Shimodaira-Hasegawa test with 1000 resamples. Trees were viewed and annotated using iTOL v.5 (Letunic and Bork, 2021).

### 2.14 Phylogenetic analysis of astacin-like metalloproteinases

For phylogenetic analysis, thirty-eight *O. ostertagi* astacin-like metalloproteinases identified in the *O. ostertagi* ES-thiol fraction were analysed in a framework of M12A domain proteins from clade V parasitic nematodes. This included three astacin-like metalloproteinases from *O. ostertagi* (CAD19995; CAD28559 and CAD11605) and one from *T. circumcincta* (Tci-MEP-1, CCR26658). Further astacin-like metalloproteinases were retrieved from the proteomes of *H. contortus* (12) (bioproject: PRJEB506; 120 proteins); *T. circumcincta* (bioproject: PRJNA72569; 244 proteins) and *H. polygyrus* (bioproject: PRJEB15396; 78 proteins) based on the presence of Interpro peptidase M12A domain (IPR001506) using an E-value threshold of <10-3. The retrieved protein set contained a total of 484 astacin-like metalloproteinases, the full list is available in Supplementary File 2. For sequence alignment we used a astacin-like metalloproteinase from *T. circumcincta* (Tci-MEP-1, CCR26658) that had been trimmed to its peptidase M12A domain (amino acids 142 – 339) to extract an alignment of these domains from the protein set. Protein domains were extracted, aligned and phylogenies reconstructed using the same pipeline for phylogenetic analysis of CAP domain proteins.

## 3. Results

### 3.1 Vaccine Trial

To confirm the protective effects of *O. ostertagi* ES-thiol-based vaccines and consistency of the effect, four independent vaccination trials were performed. Calves were immunised via the intramuscular or subcutaneous route with either *O. ostertagi* ES-Thiol adjuvanted with a saponin-based vaccine adjuvant or a saponin-based vaccine adjuvant alone, and subsequently challenged with either a trickle (Trials 1 and 2) or bolus infection (Trials 3 and 4) of *O. ostertagi* L3. Daily and cumulative FEC data from immunised and adjuvant-only control groups are shown in Figure 1. Across all four trials, immunisation with ES-thiol resulted in a significant reduction in daily and cumulative FEC relative to the adjuvant only control group (p < 0.0001 for both daily and cumulative FEC). The reductions in mean cumulative FEC for Trial 1, 2, 3 and 4 were 64%, 71%, 60% and 47%, respectively (Figure 1b). No reduction in worm burden was observed in ES-Thiol vaccinated calves in Trials 1, 2 and 3, however, a significant 26% reduction in mean worm burden in vaccinates relative to control animals was observed in Trial 4 (Supplementary Figure 1). Furthermore, ES-Thiol vaccination resulted in a significant increase in serum levels of ES-Thiol specific IgG1 and IgG2 in all four trials (Supplementary Figure 2).

**Figure 1.**
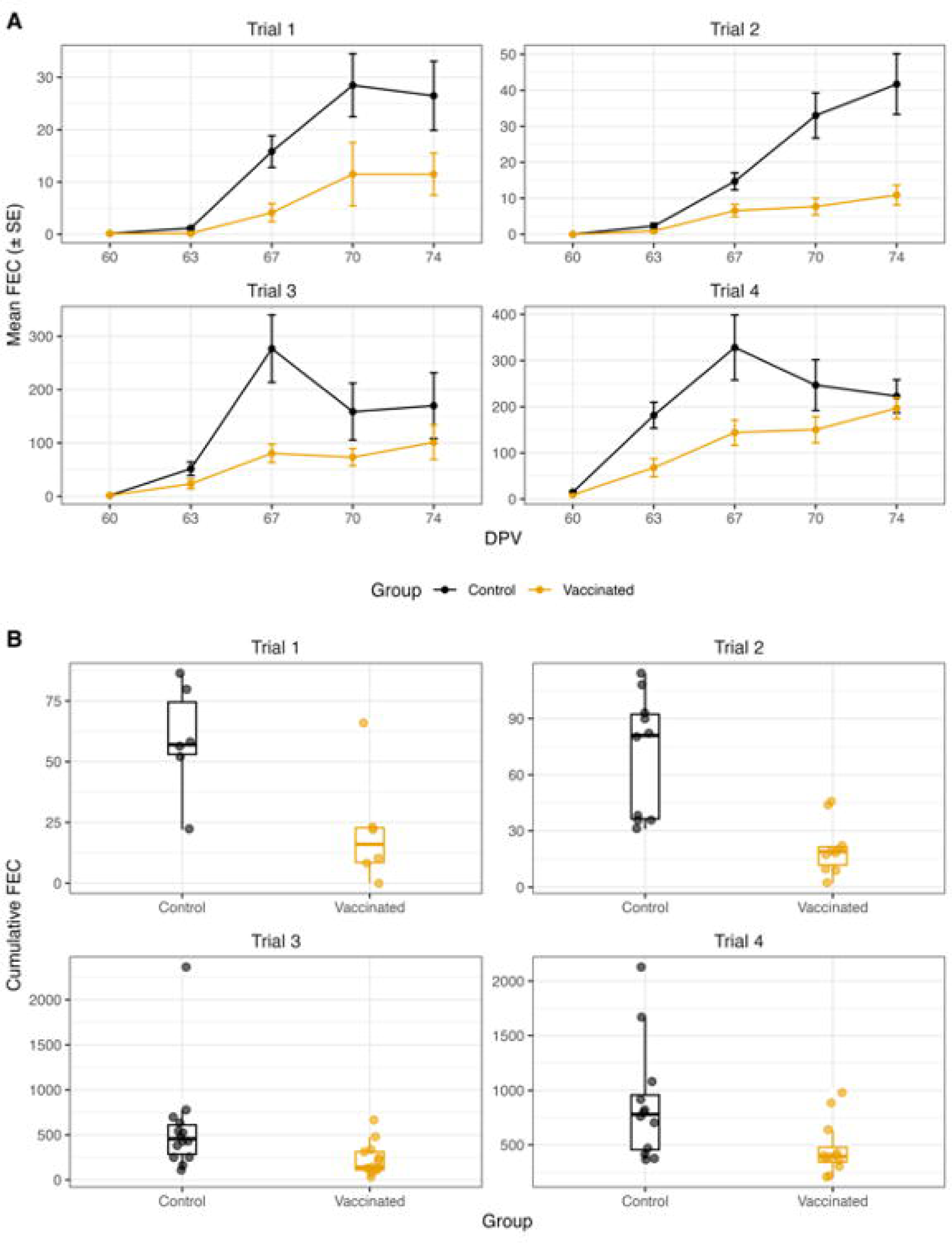
*O. ostertagi* ES-thiol vaccine trial data. (A) Daily mean faecal egg counts over time ± standard errors or the mean for Trials 1-4. (B) Boxplots of cumulative faecal egg count (cFEC) data for Trials 1-4. Individual data points represent cFEC calculated for each individual calf and boxplots indicate the median cFEC and interquartile range. Control = calves vaccinated with a saponin-based vaccine adjuvant alone. Vaccinated = calves vaccinated with ES-thiol plus a saponin-based vaccine adjuvant.

### 3.2 Generation of an *O. ostertagi* protein database

To generate a sequence database against which to search proteomic spectra data, we used Pac-Bio isoform sequencing (Iso-Seq) of *O. ostertagi* mRNA isolated from adult male and female parasites. This resulted in the generation of 15,521 and 13,387 unique open reading frames (ORFs) for male and female parasites, respectively (Figure 2). For each dataset, the average read length was 1300 bp (males) and 1459 bp (females). Completeness of each transcriptome database was assessed using Benchmarking Universal Single-Copy Orthologs (BUSCO) analysis using the metazoa lineage as the reference dataset. We identified 352 of 954 (36.9 %) and 420 of 954 (44.0 %) metazoan BUSCOs for the male and female transcript datasets, respectively, which is likely due to the relatively low depth of sequencing in Iso-seq.

**Figure 2.**
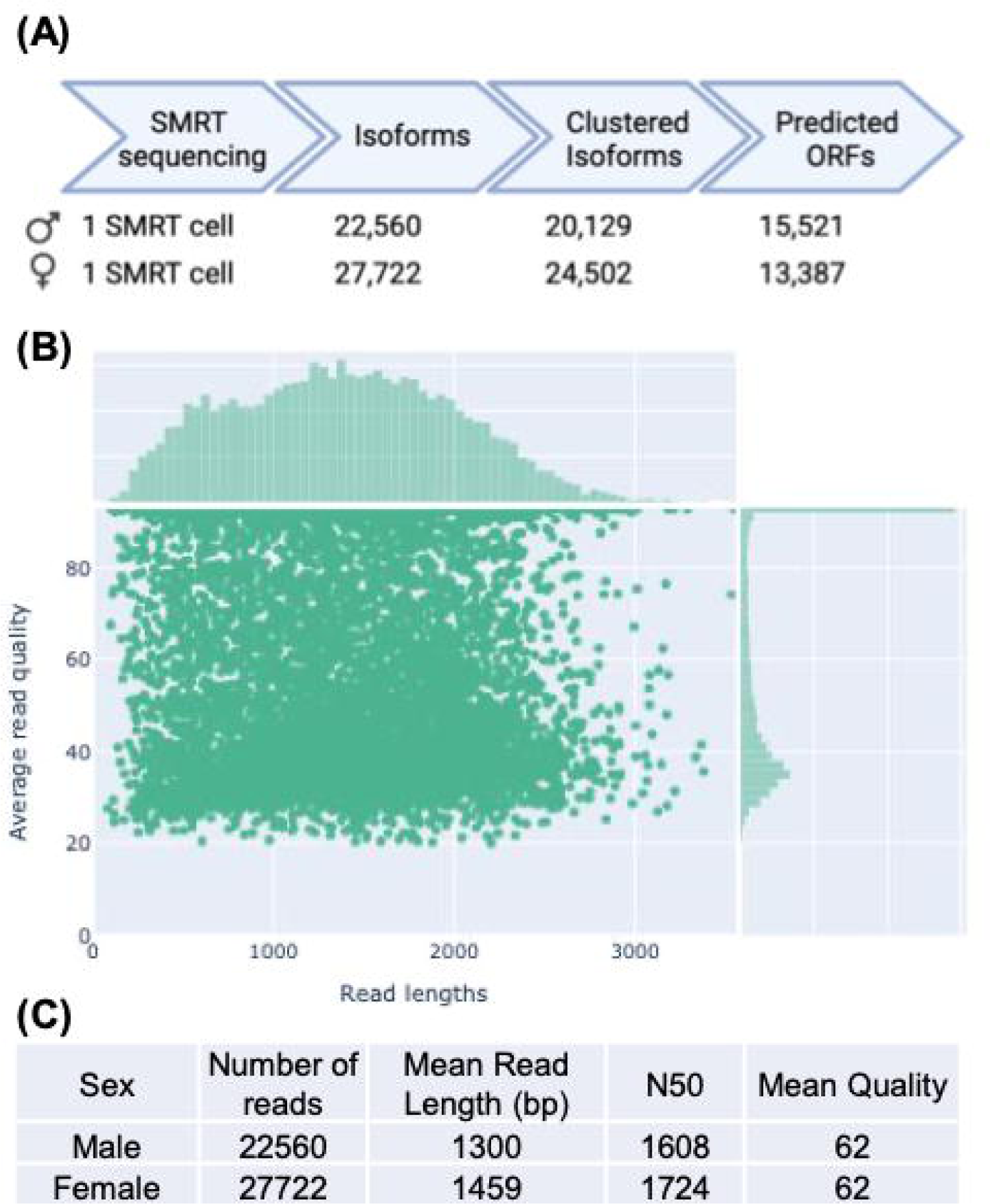
Summary of PacBio Iso-Seq sequencing for adult *O. ostertagi* males and females. (A) Bioinformatic pipeline and number of full-length isoforms generated for *O. ostertagi* male and female parasites. (B) Histogram showing read length (bp, x-axis) against read quality (Phred score, y-axis) for all male and female isoforms (50,282 isoforms in total). (C) Sequencing summary statistics for *O. ostertagi* male and female Iso-Seq libraries.

Finally, to produce a non-redundant dataset that represented both male and female *O. ostertagi* encoded proteins, protein sequences derived from both datasets were merged and redundant sequences removed by clustering using CD-HIT, based on an identity threshold of 100% across the length of each protein. This resulted in an *O. ostertagi* adult male and female database containing 22,698 unique proteins. Raw PacBio Iso-Seq data generated in this study have been deposited in the Sequence Read Archive (SRA) (https://www.ncbi.nlm.nih.gov/sra) with BioProject ID: PRJNA898386.

### 3.3 Preparation of *O. ostertagi* ES-thiol proteins

*O. ostertagi* ES-thiol proteins were purified from *O. ostertagi* adult ESPs by affinity purification. For production of sufficient material for small scale vaccination studies, 11 mg of *O. ostertagi* ESP was used as starting material, from which 0.9 mg were recovered after thiol-affinity chromatography, representing a 8.2 % protein recovery. SDS-PAGE analysis of *O. ostertagi* ES and ES-thiol is shown in Supplementary Figure 3. In both protein fractions the range of stained proteins was similar (approx. 3 – 98 kDa.), however the pattern of stained proteins differed between the two preparations. In particular, proteins at approx. 30 kDa, 45 kDa and 48 kDa in the ES-thiol fraction were enriched relative to unfractionated ES (Supplementary Figure 3). For vaccination trials presented in this study, along with the proteomic characterisation of *O. ostertagi* ES-thiol, four different batches of *O. ostertagi* ES-thiol were used. Each batch was analysed by SDS-PAGE and no visible differences in Coomassie stained protein profiles were observed. The ES-thiol preparation used in vaccine Trial 3 was subsequently subjected to proteomic analysis detailed in Section 3.4 below.

### 3.4 Proteomic analysis of *O. ostertagi* ES-thiol

To identify *O. ostertagi* ES-thiol proteins we used two complementary approaches; a proteomic analysis of tryptic peptides from a Coomassie stained SDS-PAGE gel and a proteomic analysis of tryptic peptides from an *in vitro* liquid trypsin digest. Peptides from both approaches were analysed by LC-MS/MS and resulted in the identification of 490 unique proteins in *O. ostertagi* ES-thiol (Supplementary Table 2).

Of the 490 *O. ostertagi* ES-thiol proteins identified, the majority (476, 97.1%) had blast hits against proteins from other parasitic nematodes. The most frequent top hits were against *T. circumcincta* proteins (226 hits) and *Haemonchus contortus* proteins (110 hits). A total of 14 sequences (2.9%) encoded novel proteins with no database match (Supplementary Table 2). When examined for the presence of a secretion peptide, 246 proteins (50.2%) contained a predicted signal peptide (Supplementary Table 2 and Figure 3A), while 12 proteins (2.4%) contained one or more transmembrane domains (Supplementary Table 2).

**Figure 3.**
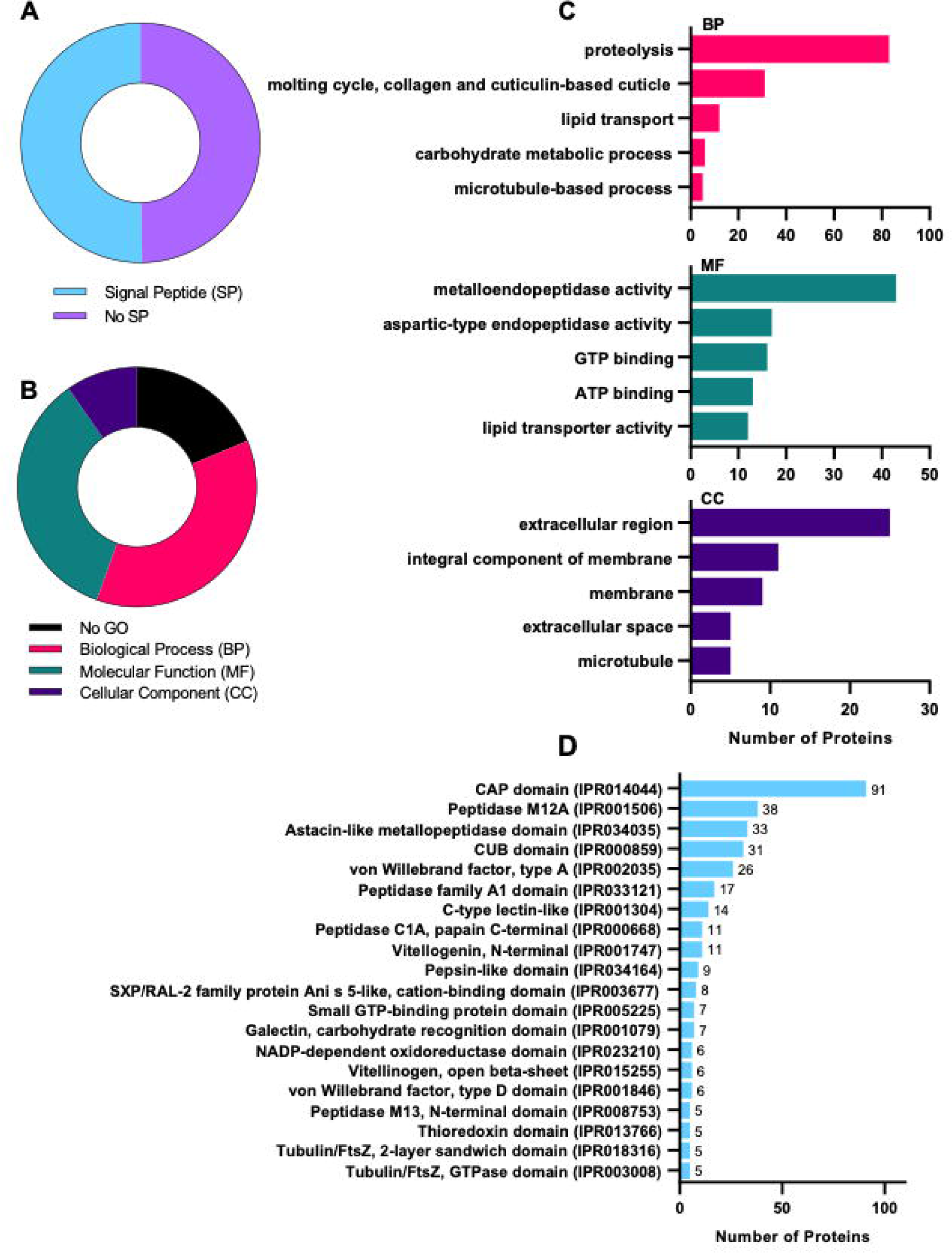
GO classification and Interpro domain classification of identified *O. ostertagi* ES proteins. (A) Proportion of ES proteins containing a secretion signal (n=490). (B) GO distribution of identified ES proteins. Proteins are classified into three broad GO categories: biological process (red); molecular function (green) and cellular component (purple). (C) Direct GO count showing the top five GO terms for each category. Some proteins are included in more than one category. (D) Interpro domain classification of identified *O. ostertagi* ES proteins showing top twenty categories. Some proteins are included in more than one category.

All identified ES-thiol proteins were classified by gene ontology (GO) analysis and assigned to categories, which included biological processes (BP), molecular function (MF) and cellular compartment (CC) (Figure 3, panel B and C). Interestingly, 182/490 (37.1%) proteins did not return GO hits in either BP, MF or CC categories. This lack of GO annotation highlights a significant knowledge gap surrounding secreted proteins produced by parasitic helminths. Of the proteins with an associated GO term, within the BP category, the top GO term was “proteolysis”, which contained 83 unique proteins (Figure 3, panel C). Proteinases were also represented in the MF classification, with “metalloendopeptidase activity” and “aspartic endopeptidase activity” containing 43 and 17 unique proteins, respectively (Figure 3, panel C). Within CC category, the top GO terms were “extracellular region” and “integral membrane protein”, with 25 and 11 proteins, respectively (Figure 3, panel C).

To get a deeper insight into protein function, the 490 *O. ostertagi* ES-thiol proteins were analysed using Interpro domain classification (Figure 3, panel D). The most numerous protein family in the *O. ostertagi* ES-thiol preparation was the SCP/TAPS family and accounted for 91/490 proteins, containing either a single or double Interpro CAP domain (IPR014044) (Figure 3, panel D). The top-20 most numerous protein families are shown in Figure 3, panel D. Within the top-20 most frequently observed domains, proteases were highly represented, including domains from: metalloproteinases (peptidase M13, N-terminal domain [IPR008753]; peptidase M12A [IPR001506]); aspartyl proteases (Peptidase family A1 domain [IPR033121]; pepsin-like domain [IPR034164]; astacin-like metallopeptidase domain [IPR034035]) and cysteine proteinases: (peptidase C1A, papain C-terminal [IPR000668]) (Figure 3, panel D).

### 3.5 Phylogenetic analysis of *O. ostertagi* CAP domain proteins

The proteomic analysis of *O. ostertagi* ES-thiol demonstrated that there is a large number of CAP domain proteins within this protein fraction. In addition, previous studies has shown that Oo-ASP-1 and Oo-ASP-2 (both CAP domain proteins) are the most abundant proteins within *O. ostertagi* ES-thiol (Geldhof et al., 2003a; Meyvis et al., 2007a). Thus, given the importance of CAP domain proteins in the *O. ostertagi* ES-thiol fraction we performed a comprehensive phylogenetic analysis of this protein family.

Within the *O. ostertagi* ES-thiol fraction we identified 91 CAP domain containing proteins which match the InterPro CAP domain (IPR014044) (Figure 4 and Supplementary Table 3). The majority of *O. ostertagi* CAP domain proteins (75/91, 82.4%) have a predicted N-terminal secretion signal, with 43/91 containing a single CAP domain and 48/91 with a double CAP domain (Supplementary Table 3). Phylogenetic analysis of CAP domain proteins from *O. ostertagi* ES-thiol, along with CAP domain proteins from other parasitic nematodes is shown in Figure 4. Within this phylogeny, previously characterised *O. ostertagi* ASP vaccine candidates [ASP-1 (Vercauteren et al., 2003) and ASP-2 (Geldhof et al., 2003a)] and other *O. ostertagi* ASPs [Oo-ASP-3 and Oo-ASP-4 (Visser et al., 2008)] are highlighted. In addition, we have highlighted nine ASPs from *Ancylostoma caninum*, which have become the canonical set of hookworm ASPs (these include: Acan-NIF (Moyle et al., 1994); Acan-PI (Del Valle et al., 2003); Acan-ASP-1 (Hawdon et al., 1996); Acan-ASP-2 (Hawdon et al., 1999); Acan-ASP-3, Acan-ASP-4, Acan-ASP-5, Acan-ASP-6 from (Zhan et al., 2003) and Acan-ASP-7 (Datu et al., 2008)] (Figure 4).

**Figure 4.**
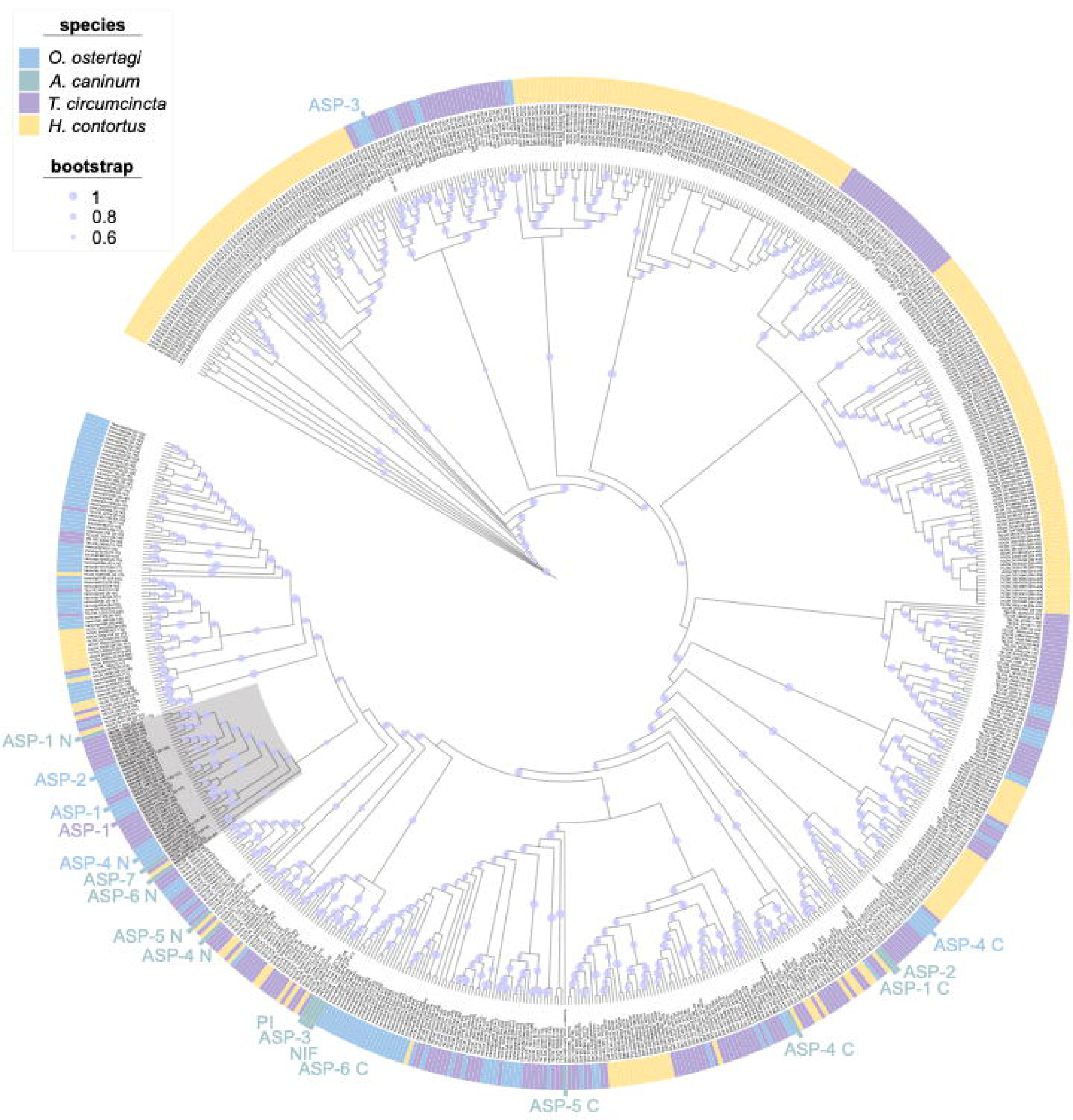
Maximum likelihood phylogeny of CAP domain proteins identified in Oost ES-thiol and other parasitic nematodes. Due to the structure of CAP domain proteins, which contain either a single or double domain, only the CAP protein domain was used for phylogenetic reconstruction. CAP domain proteins from *O. ostertagi* (blue), *T. circumcincta* (purple), *A. caninum* (green) and *H. contortus* (yellow) were included in the analysis. Accession numbers for each identified CAP protein are shown in the tree and the coordinates of the CAP domain that was used for phylogenetic reconstruction is shown in square brackets. Known ASPs are highlighted and the N– and C– terminal CAP domains from two-domain ASPs are noted as “N” or “C”. A subtree (38 nodes) that contains *O. ostertagi* ASP-1 and *T. circumcincta* ASP-1 is highlighted grey. Confidence values >0.6 are shown on branches of the tree.

The phylogenetic reconstruction shows several well supported clades that contain large numbers of homologous CAP domain proteins from the same nematode genus, which indicates recent duplication and diversification (Figure 4). Within the phylogeny, Oo-ASP-1 and Oo-ASP2 (both single domain CAP proteins) resolve together in a well-supported clade (Figure 4). The phylogeny demonstrates that Oo-ASP-1 (CAD23183) is closely related to *T. circumcincta* ASP-1 (CBJ15404) plus 3 other identified proteins in *O. ostertagi* ES-thiol, represented by transcripts including: transcript/16135; transcript/14620 and transcript/15497 (Figure 4). In contrast, for OoASP-2 (CAD56659) there is no closely related *T. circumcincta* orthologue, whereas there are six related proteins in *O. ostertagi* ES-thiol, represented by transcripts: transcript/15383; transcript/15960; transcript/18805; transcript/14958; transcript/15874 and transcript/15062 (Figure 4).

### 3.6 Phylogenetic analysis of *O. ostertagi* Astacin-like Metallopeptidase

In our proteomic analysis, the second most numerous family of proteins in *O. ostertagi* ES-thiol were the astacin-like metalloproteinases, with hits to peptidase M12A domain (IPR001506, 38 proteins) and astacin-like metallopeptidase domain (IPR034035, 33 proteins) (Figure 5). Prior to this analysis, three *O. ostertagi* astacin-like metalloproteinases had been identified in *O. ostertagi* ES (including CAD19995; CAD28559 and CAD11605) by immunoscreening with antibody from immune cattle (De Maere et al., 2002). In addition, we have previously shown that an astacin-like proteinase from *T. circumcincta* (Tci-MEP-1, CCR26658) is present in *T. circumcincta* ES material of infective stage parasites, and recognised by both IgA and IgG responses of immune sheep (Nisbet et al., 2019; Smith et al., 2009). In addition Tci-MEP-1 is a component of a recombinant cocktail vaccine that protects sheep against *T. circumcincta* infection (Nisbet et al., 2013). Thus, given that astacin-like metallopeptidases are abundant in the protective native *O. ostertagi* ES-thiol vaccine, and are a protective antigen in *T. circumcincta* vaccine, we investigated this family further in *O. ostertagi*.

**Figure 5.**
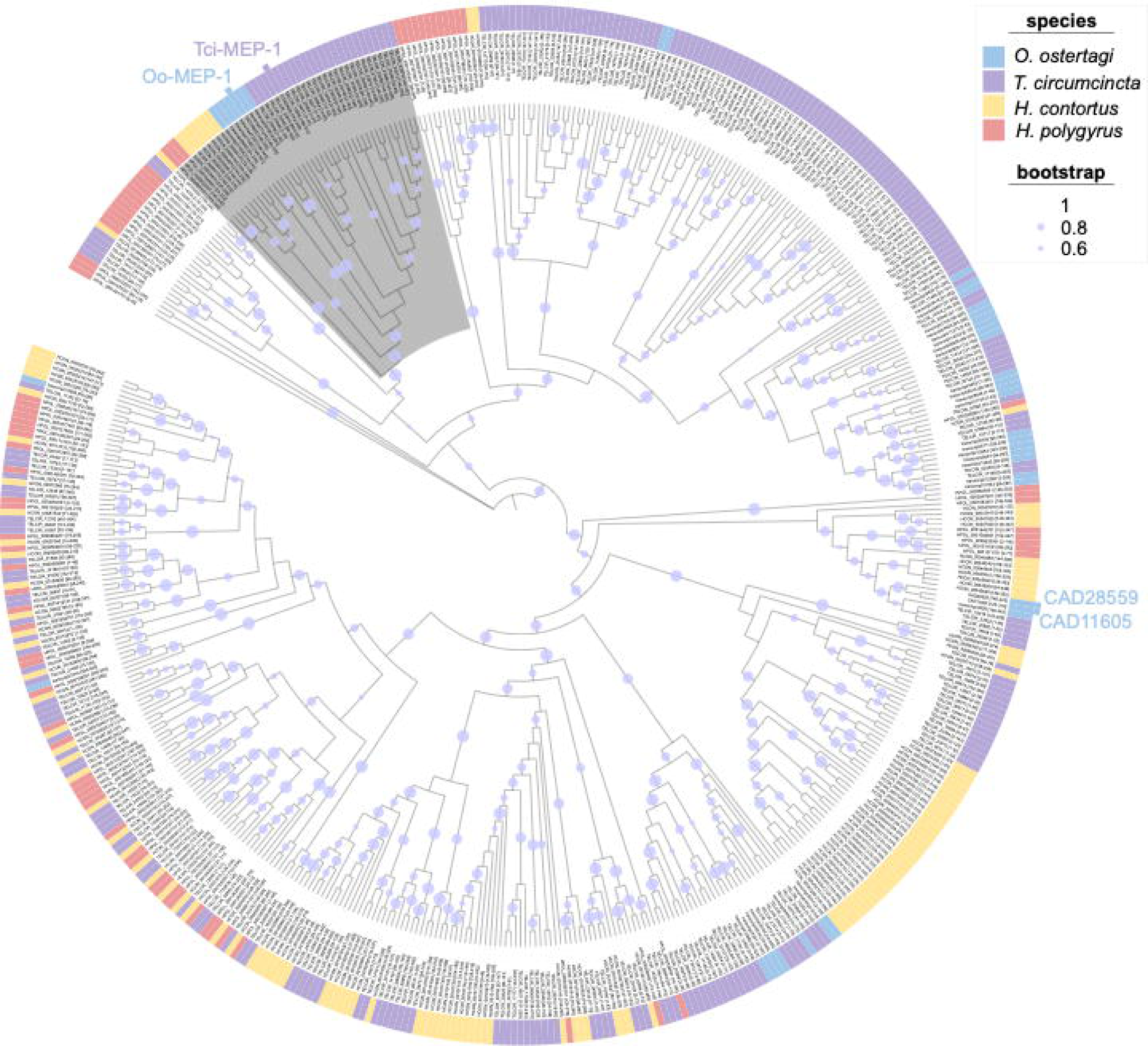
Maximum likelihood phylogeny of astacin-like metalloproteinases in Oost ES-thiol and other parasitic nematodes. Astacin-like peptidase M12A domain proteins from *O. ostertagi* (blue), *T. circumcincta* (purple), *H. contortus* (yellow) and *H. polygyrus* (pink) were included in the analysis. Accession numbers for each astacin-like metalloproteinase are shown in the tree and the coordinates of the peptidase M12A domain that was used for phylogenetic reconstruction is shown in square brackets. Previously characterised astacin-like proteinases from *O. ostertagi* and *T. circumcincta* are highlighted. A subtree (43 nodes) that contains *O. ostertagi* MEP-1 and *T. circumcincta* MEP-1 is highlighted grey. Confidence values >0.6 are shown on branches of the tree.

Within the *O. ostertagi* ES-thiol fraction there were 38 putative astacin-like metalloproteinases which match the InterPro Peptidase M12A domain (IPR001506) (Figure 5 and Supplementary Table 2 and 4). Relatively few of the *O. ostertagi* astacin-like metalloproteinases (9/38, 23.7%) had a predicted N-terminal secretion signal, and a single protein had a predicted transmembrane domain (Supplementary Table 2 and 4). The identified astacin-like metalloproteinases had a multi-domain structure that is common within this family of proteinases, with 31 containing a CUB domain (IPR000859) and a single proteinase with a ShKT domain (IPR003582) (Möhrlen et al., 2003). Phylogenetic analysis of astacins from *O. ostertagi* ES-thiol, along with astacins from other parasitic nematodes is shown in Figure 5. The phylogenetic analysis shows several well supported clades that contain large numbers of homologous astacin-like metalloproteinases from the same nematode genus, which indicates recent duplication and diversification (Figure 5). Within the phylogeny, the *O. ostertagi* astacins group most closely with those from *T. circumcincta*. Of the previously described *O. ostertagi* astacins, metalloproteinase I (CAD19995) is most closely related to *T. circumcincta* MEP-1 (CAD19995) along with six astacins within the same clade (including those encoded by transcript/9188; transcript/11276; transcript/7232; transcript/11922; transcript/9025 and transcript/11256) (Figure 5).

### 3.7 Other vaccine candidates in *O. ostertagi* ES-thiol

We have previously generated a recombinant subunit vaccine against *T. circumcincta* that protects sheep from *T. circumcincta* infection (Nisbet et al., 2013). As *T. circumcincta* and *O. ostertagi* are both clade V nematodes, and phylogenetically closely related, we searched *O. ostertagi* ES-thiol proteins for orthologues of *T. circumcincta* vaccine antigens (Table 1). The *T. circumcincta* recombinant vaccine contains eight proteins, including: Tci-SAA-1; Tci-MIF-1; Tci-ASP-1; Tci-TGH-2; Tci-CF-1; Tci-ES20; Tci-MEP-1 and Tci-APY-1 (for accession numbers and description of proteins see Table 1). Of these proteins, homologues of Tci-ASP-1, Tci-CF-1, Tci-ES20, Tci-MEP-1 and Tci-APY-1, which meet the criteria of reciprocal best blast-hits, were detected in *O. ostertagi* ES-thiol proteins (Table 1).

**Table 1.**
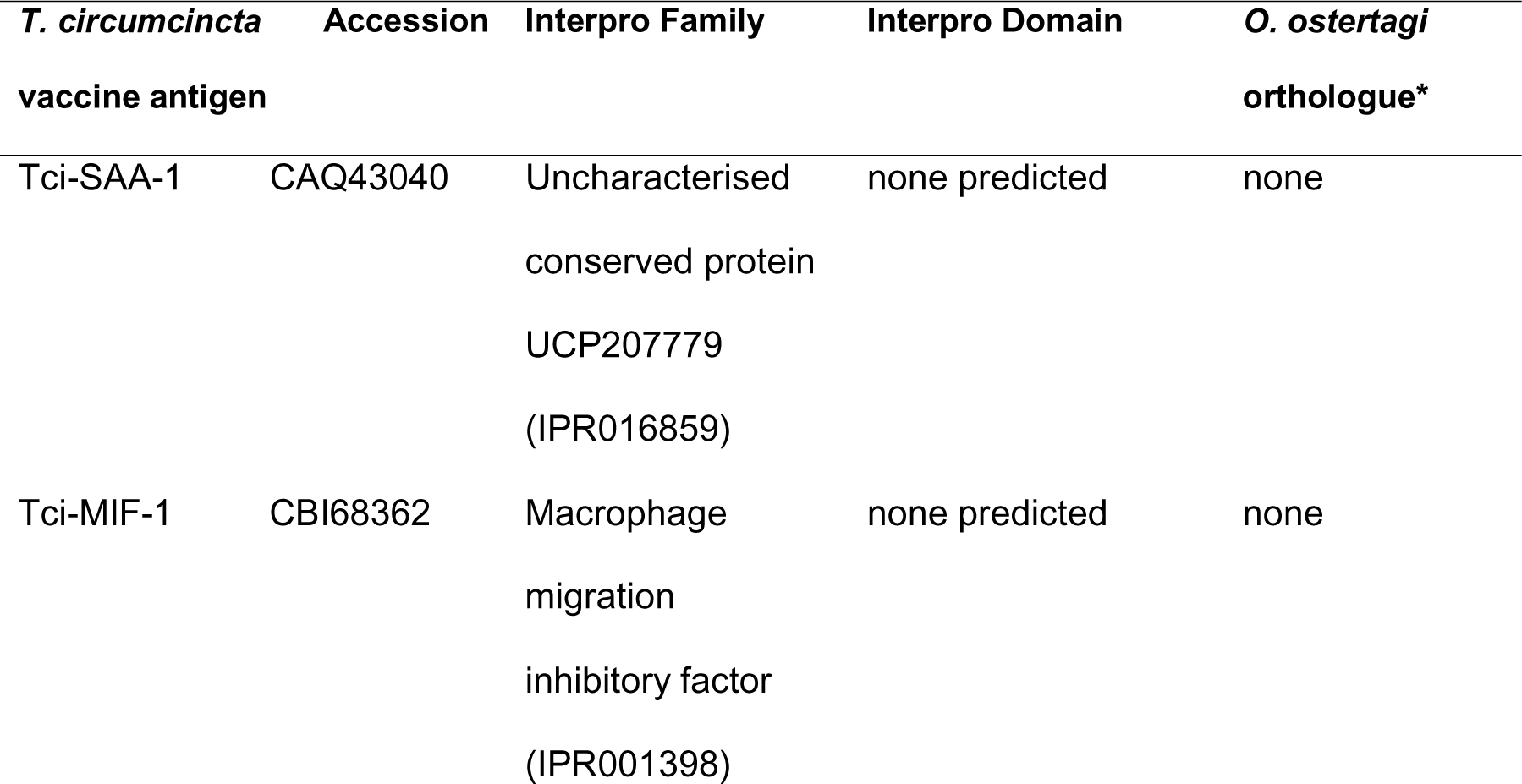
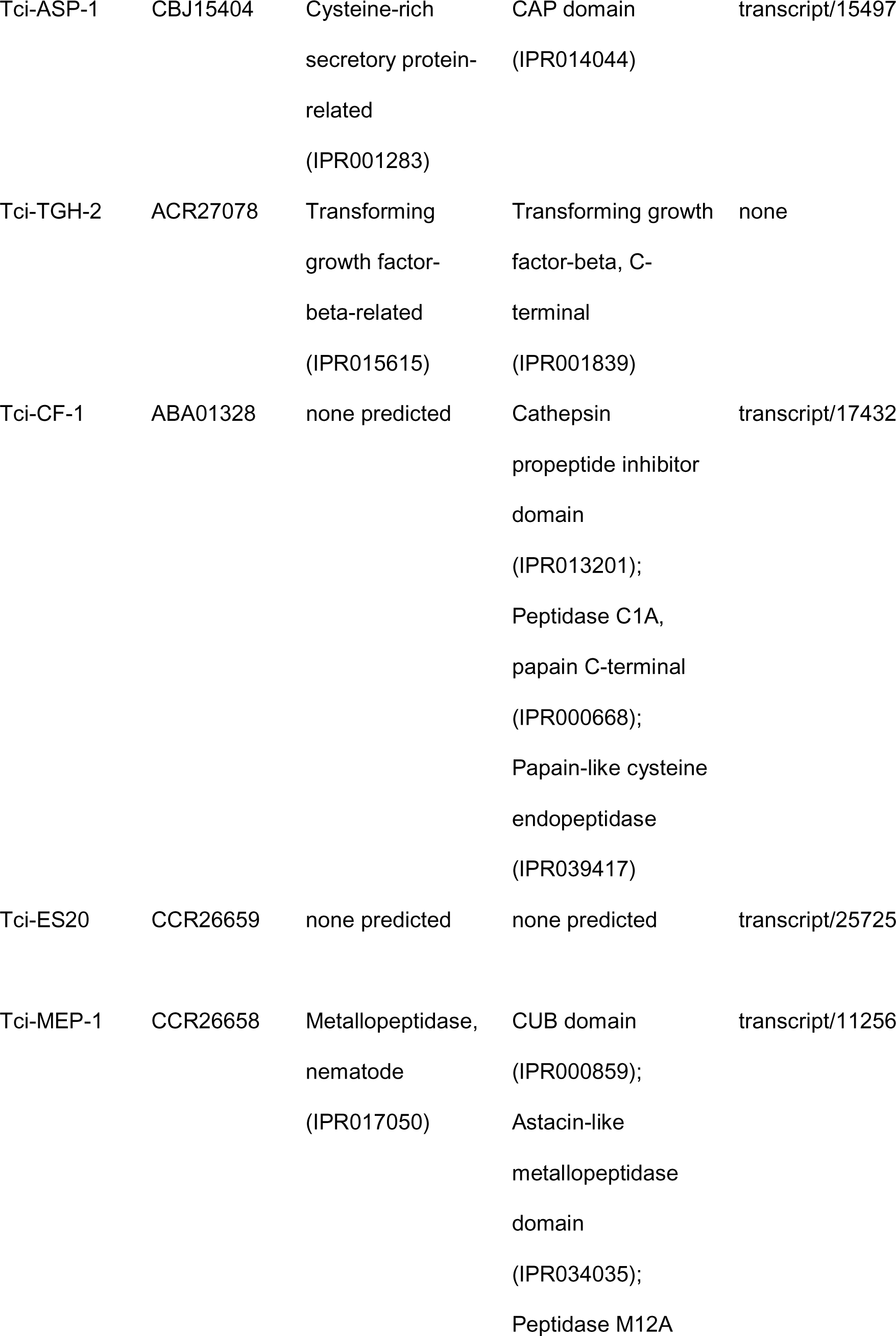
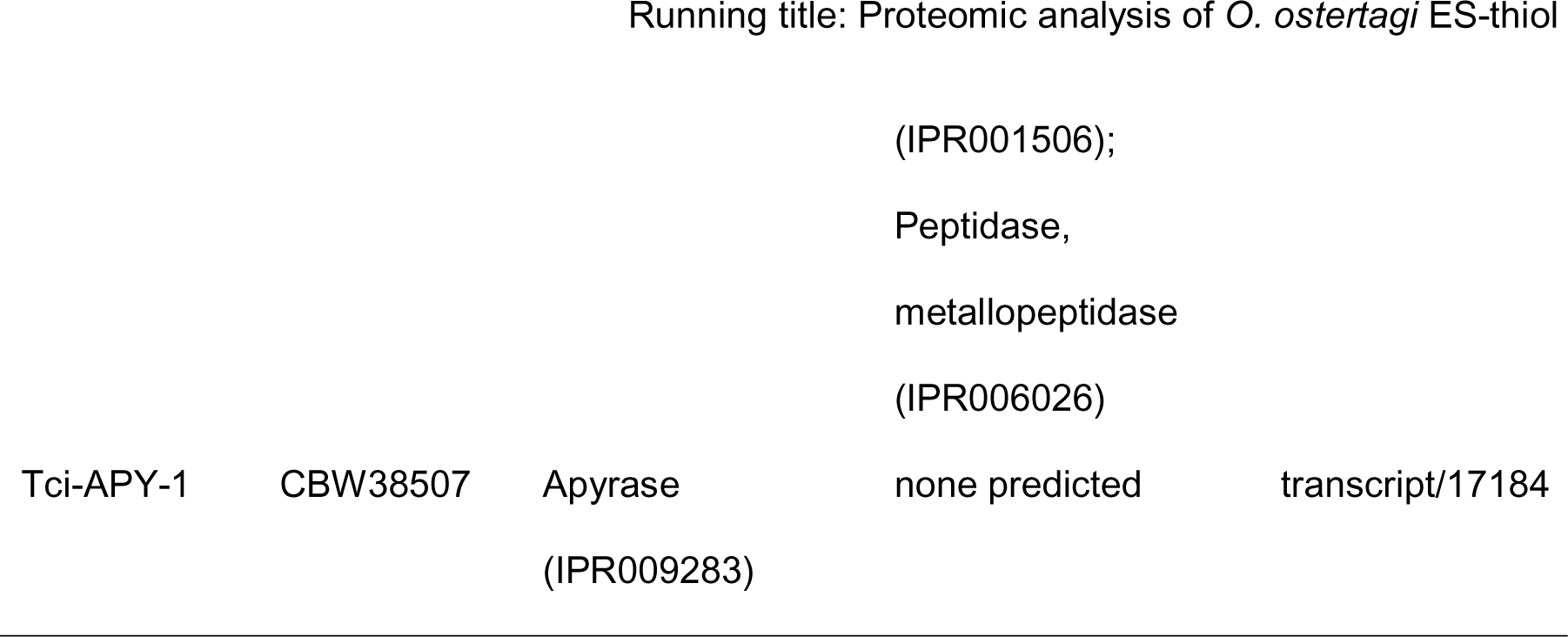
Identification of *T. circumcincta* vaccine orthologs in *O. ostertagi* ES-thiol, based on reciprocal blast best hits.

## 4. Discussion

Our current study confirms that antigens derived from *O. ostertagi* ES-thiol protect cattle against *O. ostertagi* infection when used as a vaccine. In agreement with previous reports, the main protective effect is a significant reduction in cumulative faecal egg counts, while adult worm burdens between vaccinates and controls were not significantly different in three (Trial 1; Trial 2 and Trial 3) of the four trials (Geldhof et al., 2004, 2003a, 2002; Meyvis et al., 2007a). In addition, our study demonstrates that protection with the vaccine is reproducible, with significant levels of protection across four independent vaccination-challenge experiments.

The commercial success of vaccines depends not only on robust and reproducible levels of protection, but the ability to produce the vaccine that is economically viable to use. There are currently two commercial vaccines, based on native parasite antigens, which meet these criteria. The first, commercially known as Bovilis® Huskvac (produced by MSD animal health), is a live attenuated vaccine that protects against the cattle lungworm, *Dictyocaulus viviparus* (Jarrett et al., 1960). The second, commercially known as Barbervax®, protects sheep against infection by the barber pole worm, *H. contortus*, and is based upon native parasite antigens isolated by lectin affinity chromatography (Nisbet et al., 2016; Smith et al., 1994). Although the *O. ostertagi* ES-thiol vaccine offers high levels of protection against *O. ostertagi* infection, its costly and time-consuming production negates its use as a commercial product. The high cost associated with this vaccine result from complexities of its production, which includes: isolation of adult parasites from infected cattle; *in vitro* culture of the parasite to generate ESPs and recovery of low amounts of protective ES-thiol antigens from ESPs. For these reasons, commercial exploitation of this vaccine is not feasible.

The aim of this study was to characterise the proteome of *O. ostertagi* ES-thiol using high-resolution shotgun proteomics to inform future vaccine antigen discovery projects, with the capacity to complement the current best, ASP-based, vaccine candidates (Geldhof et al., 2003a, 2002; Meyvis et al., 2007a). Previous proteomic analysis of the ES-thiol, and sub-fractions of the ES-thiol, have demonstrated that the single domain ASPs Oo-ASP-1 and Oo-ASP-2 are the principal protective antigens (Geldhof et al., 2003a; Meyvis et al., 2007a).

However, in addition to Oo-ASP-1 and Oo-ASP-2, it is likely there are other protective antigens within this protein preparation. This was demonstrated by an *O. ostertagi* vaccine trial using three sub-fractions of ES-thiol, which included: an ASP-enriched fraction; a cysteine proteinase (CP) enriched fraction and a “remaining proteins” fraction (Meyvis et al., 2007a). Following *O. ostertagi* challenge, all three fractions were protective and resulted in significantly reduced cumulative FEC relative to control animals (Meyvis et al., 2007a). Importantly, it was shown that antibodies from animals vaccinated with the CP enriched fraction and the “remaining proteins” fraction did not bind antigens from the ASP enriched fraction. Thus, demonstrating that in addition to the ASPs, there are other protective antigens within ES-thiol (Meyvis et al., 2007a).

To conduct a comprehensive proteomic assessment of *O. ostertagi* ES-thiol, we first needed a protein sequence database, against which proteomic data could be searched. As there is currently no available *O. ostertagi* genome, we used isoform sequencing (Iso-Seq) to generate a library of full-length transcripts from male and female *O. ostertagi* parasites. Using this approach, we generated 15,521 male and 13,387 female high quality unique full-length transcripts, significantly increasing the number of *O. ostertagi* sequences available in the NCBI database. With the database in place, we used high-resolution shotgun proteomics to identify 490 proteins within *O. ostertagi* ES-thiol. Our analysis significantly extends previous proteomic characterisation of *O. ostertagi* ES-thiol (Geldhof et al., 2003a), and demonstrates that the fraction is relatively complex. Importantly, from the proteins that we identified, previously described protective antigens Oo-ASP-1 and Oo-ASP-2 (Geldhof et al., 2003a; Meyvis et al., 2007a) as well as cysteine proteinases (Geldhof et al., 2002) were present. Of the identified proteins approx. 50% contained signal peptide, indicating these are likely secreted via the classical secretory pathway. The remaining proteins lack a signal peptide, but were secreted or excreted nonetheless. It is possible that these proteins are packaged for secretion via alternative pathways. Such pathways include direct transfer across the plasma membrane and packaging of protein cargo within plasma membrane derived extracellular vesicles (EVs) (Cohen et al., 2020). It has been shown that many secreted parasite proteins, especially those contained within EVs, lack a classical N-terminal secretion signal (Silverman et al., 2008). The cellular mechanism of protein secretion in parasitic nematodes, and biological significance of partitioning proteins across different secretory pathways awaits further investigation.

In terms of protein abundance within *O. ostertagi* ES-thiol, it has been shown previously that Oo-ASP-1 and Oo-ASP-2 are the dominant CAP proteins (Geldhof et al., 2003a). Our analysis shows that in addition to Oo-ASP-1 and Oo-ASP-2 there is a large diversity of CAP domain proteins within this protein preparation (Supplementary Table 2 and 3 and Figure 4). Although we can detect a large number of additional CAP-domain proteins (91 CAP domain proteins in total), their relative abundance is unknown, and therefore their contribution towards vaccine efficacy remains to be determined. It is known that CAP-domain proteins are ubiquitously present in ESPs of parasitic nematodes, and often upregulated in parasitic phases of their lifecycle (Heizer et al., 2013; Hewitson et al., 2011; International Helminth Genomes Consortium, 2019; Viney, 2017). Expression profiles and the secreted nature of *O. ostertagi* CAP domain proteins point to a role in host-parasite interactions, yet specific roles for most CAP domain proteins expressed by helminths remain unknown, with few exceptions. The hookworm CAP domain protein Na-ASP-2 from *Necator americanus*, has been shown to have immunomodulatory function, and is able to supress B cell receptor signalling and reduce neutrophil recruitment and therefore plays an important role in host-parasite interactions (Bower et al., 2008; Tribolet et al., 2015). Given the abundance of CAP domain proteins in *O. ostertagi* ES-thiol, and that they are expressed and secreted in parasitic life stages of the parasite, it is likely that they also play an important role in host-parasite interactions.

Proteases and proteinase inhibitors are also numerous in the secretomes of parasitic nematodes (Hewitson et al., 2011; International Helminth Genomes Consortium, 2019; Martín-Galiano and Sotillo, 2022; Viney, 2017) and this is also true of *O. ostertagi* ES-thiol. Our proteomic analysis reveals an abundance of proteinases in this fraction, representing three distinct classes, including: metallo-; aspartyl– and cysteine proteinases (Figure 3, panel D). In parasitic nematodes it is thought that secreted proteinases and proteinase inhibitors perform diverse tasks linked to parasitism, including immunomodulation, host tissue penetration and modification of the environment and digestion of blood (McKerrow et al., 2006). Within *O. ostertagi* ES-thiol, M12 astacin-like metalloproteinases are the most numerous family of proteinases, with 38 proteins containing a peptidase M12A domain (IPR001506) and 33 proteins with a astacin-like metallopeptidase domain (IPR034035) (Supplementary Table 2 and 4 and Figure 5). M12 astacins are one of the most abundant proteins in the secretome of parasitic nematodes (Martín-Galiano and Sotillo, 2022). This family is extensively expanded in nematode clades IVa and V, where they have been suggested to play a role in tissue migration and immune modulation (International Helminth Genomes Consortium, 2019). Given the abundance of astacins in *O. ostertagi* ES thiol, and the importance of these proteinases in other parasitic nematodes, this family is likely to play a key role in host-parasite interactions. Again, as with the other protein families described here, the physiological role of secreted proteinases, as well as the role they play in generation of protective immunity in *the O. ostertagi* ES-thiol vaccine, remains to be determined.

A commercially viable approach to produce anti-parasite vaccines is through recombinant expression of critical parasite antigens that are important for host-parasite interactions. An example of this includes the vaccine against the tapeworm *Taenia solium*, which is based on a single recombinant parasite antigen, designated TSOL18 (Flisser et al., 2004). This vaccine, commercially sold as Cysvax™, is highly efficacious, and provides 100% protection against the disease (Flisser et al., 2004; Jayashi et al., 2012). Recombinant vaccines also have potential to protect against parasitic nematodes. We have previously shown that a recombinant subunit vaccine based on worm ES proteins protects sheep against *T. circumcincta* infection. The vaccine contains recombinant versions of eight proteins and protects sheep against *T. circumcincta* infection, reducing both egg count (58 – 70%) and adult worm burden (56% – 75%) in vaccinated lambs (Nisbet et al., 2013). Of the eight antigens included in this recombinant vaccine, orthologues of six are found in *O. ostertagi*-ES-thiol (Table 1). It remains to be determined if these *O. ostertagi* orthologues are recognised by the host immune response and if they contribute towards ES-thiol vaccine efficacy.

In summary, our work demonstrates that the *O. ostertagi* ES-thiol vaccine offers consistent protection against homologous parasite challenge across four independent vaccine trials. Using high-resolution shotgun proteomics, we demonstrate that *O. ostertagi* ES-thiol is relatively complex, with 490 identified proteins, belonging to diverse protein families. Analysis of the *O. ostertagi* ES-thiol proteome reveals that CAP domain proteins and astacin-like metalloproteinases, which are highly abundant in the secretome of parasitic nematodes, are also highly abundant in the *O. ostertagi* ES-thiol fraction. Within the ES-thiol fraction, CAP-domain proteins dominate, and include previously identified protective antigens Oo-ASP-1 and Oo-ASP-2. Our proteomic analysis of ES-thiol will facilitate future *O. ostertagi* vaccine antigen discovery projects to complement existing *O. ostertagi* vaccine antigens.

## 5. Conflict of Interest

The authors declare that the research was conducted in the absence of any commercial or financial relationships that could be construed as a potential conflict of interest.

## 6. Author Contributions

PS and TMN conceived the study. All authors designed the research. DRGP, PS, TNM, KML, DA performed research. PG and JB trained PS in *O. ostertagi* ES-thiol purification. DRGP, PS, KML, DA, JPA, AJN, TMN analysed data. DRGP wrote the paper with contributions from all authors. All authors read and approved the final manuscript.

## 7. Data availability statement

Raw PacBio Iso-Seq data generated in this study have been deposited in the Sequence Read Archive (SRA) (https://www.ncbi.nlm.nih.gov/sra) with BioProject ID: PRJNA898386.

## 8. Funding

The work was supported in part by a Moredun Foundation Research fellowship awarded to DRGP. All authors receive funding from the Scottish Government Rural and Environment Science and Analytical Services Division (RESAS) and this work was partly funded from that source.

## 9. Acknowledgements

We thank the Bioservices Group at Moredun Research Institute for assistance with animal studies.

## 13. Supplementary Material

**Supplementary File 1.** CAP domain proteins used for phylogenetic analysis.

**Supplementary File 2.** Astacin-like metalloproteinases domain proteins used for phylogenetic analysis.

**Supplementary Figure 1.** Worm burden data for Trial 1 – 4.

**Supplementary Figure 2.** ELISA data for Trial 1 – 4.

**Supplementary Figure 3.** SDS-PAGE analysis of *O. ostertagi* ES and ES-thiol.

**Supplementary Table 1.** Experimental design of *O. ostertagi* ES-thiol vaccine Trials 1 – 4.

**Supplementary Table 2.** Summary of identified proteins in *O. ostertagi* ES-thiol.

**Supplementary Table 3.** Identified CAP domain proteins in *O. ostertagi* ES-thiol.

**Supplementary Table 4.** Identified astacin-like metalloproteinases in *O. ostertagi* ES-thiol.

## Notes

https://www.ncbi.nlm.nih.gov/sra

### Competing Interest Statement

The authors have declared no competing interest.

