## Supplementary Figure 1 - 3 for "Characterisation of protective vaccine antigens from the thiol-containing components of excretory/secretory material of *Ostertagia ostertagi*"

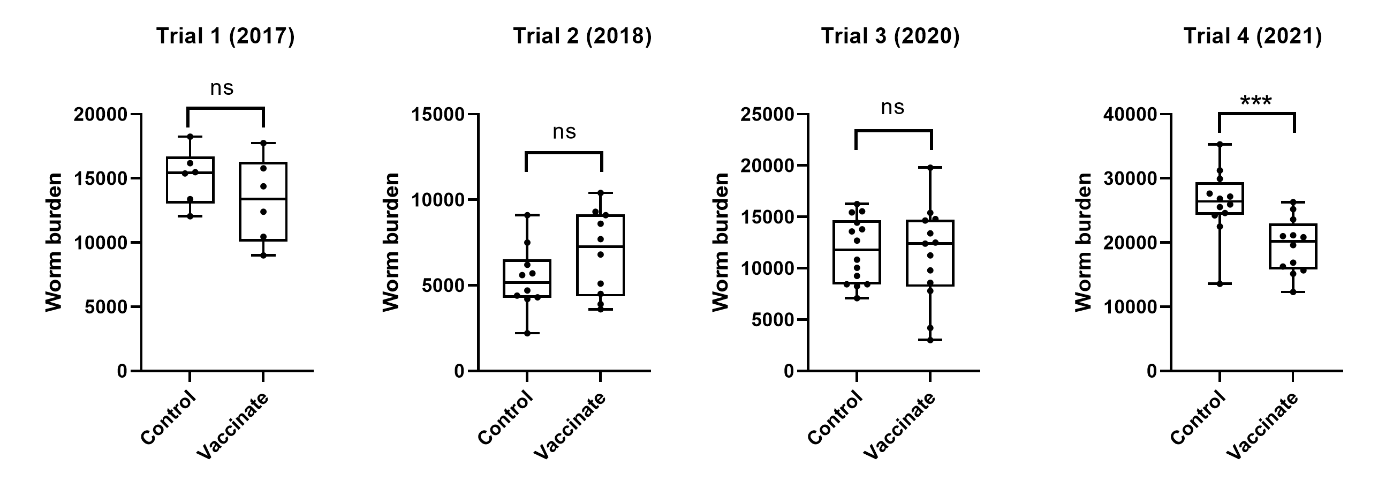

**Supplementary Figure 1.** Nematode worm burdens at post-mortem of calves vaccinated with *O. ostertagi* ES-thiol (vaccinate) or saponin-based vaccine adjuvant only (control) following challenge infection. Total worm burden (luminal and mucosal worms) at post-mortem of calves in Trial 1 (n = 6, each group), Trial 2 (n = 10, each group), Trial 3 (n = 14 for control, n = 13 for vaccinate) and Trial 4 (n = 12, each group). Individual data points represent worm burden calculated for each individual calf and boxplots indicate the median cFEC and interquartile range. Significant differences (*** p<0.001) and no significant differences (ns) between control and vaccinate groups are indicated.

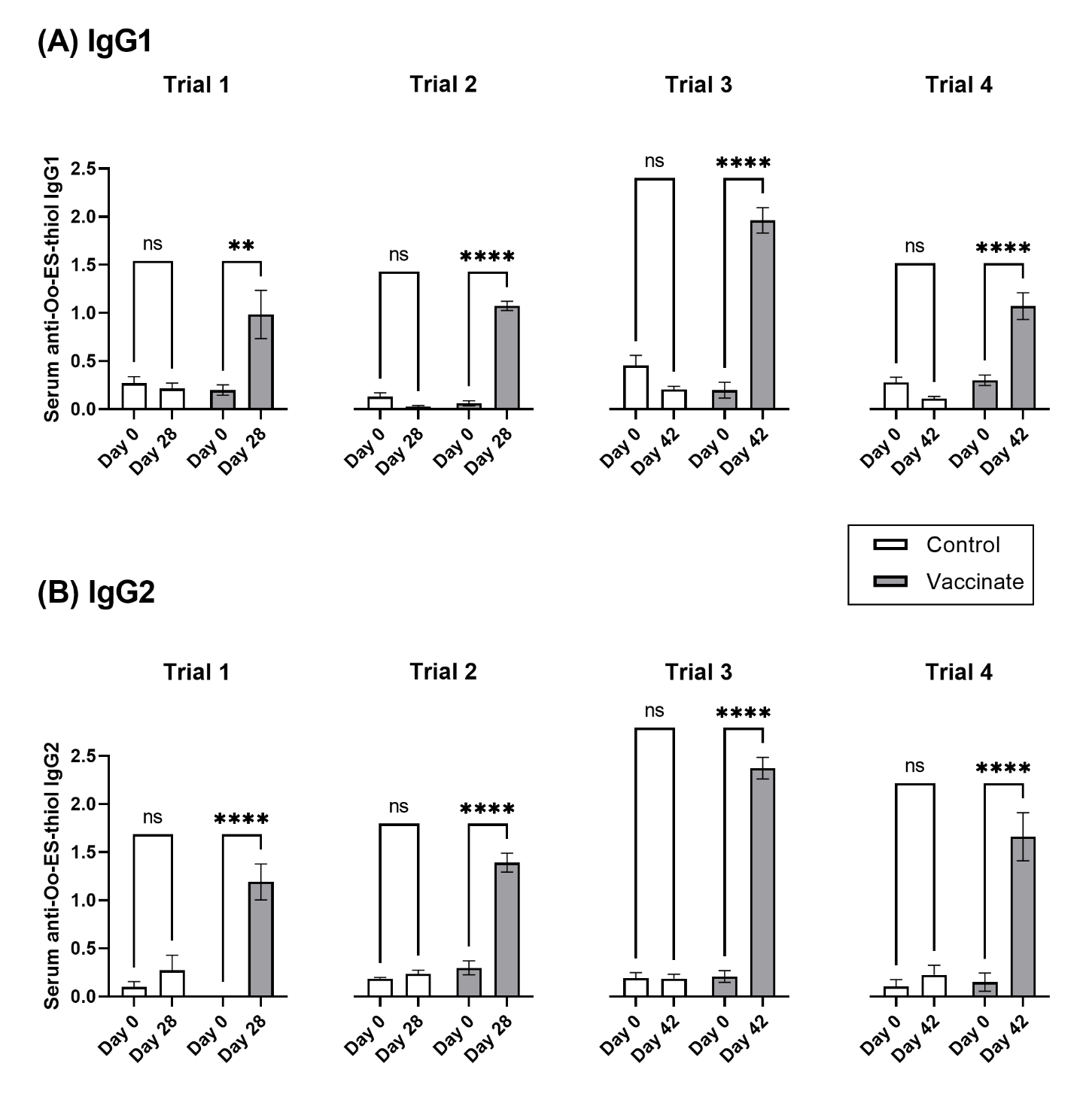

**Supplementary Figure 2.** Serum antibody responses of vaccinated cattle to *O. ostertagi* ES-thiol across four independent trials. For each Trial (Trial 1-4) cattle were immunised with either Oo-ES-thiol (vaccinate) or saponin-based vaccine adjuvant only (control). For each trial, serum IgG1 (Panel A) and IgG2 (Panel B) responses against Oo-ES-thiol were determined at day 0 (pre-vaccination) and day 28 for Trials 1 and 2 and day 42 for Trials 3 and 4. Bars show mean antibody response ± SEM (Trial 1, n = 6; Trial 2, n = 10; Trial 3, n = 14; Trial 1, n = 12). Statistically significant differences (**, p<0.01; ****, p<0.0001), and non-significant differences (ns) are indicated.

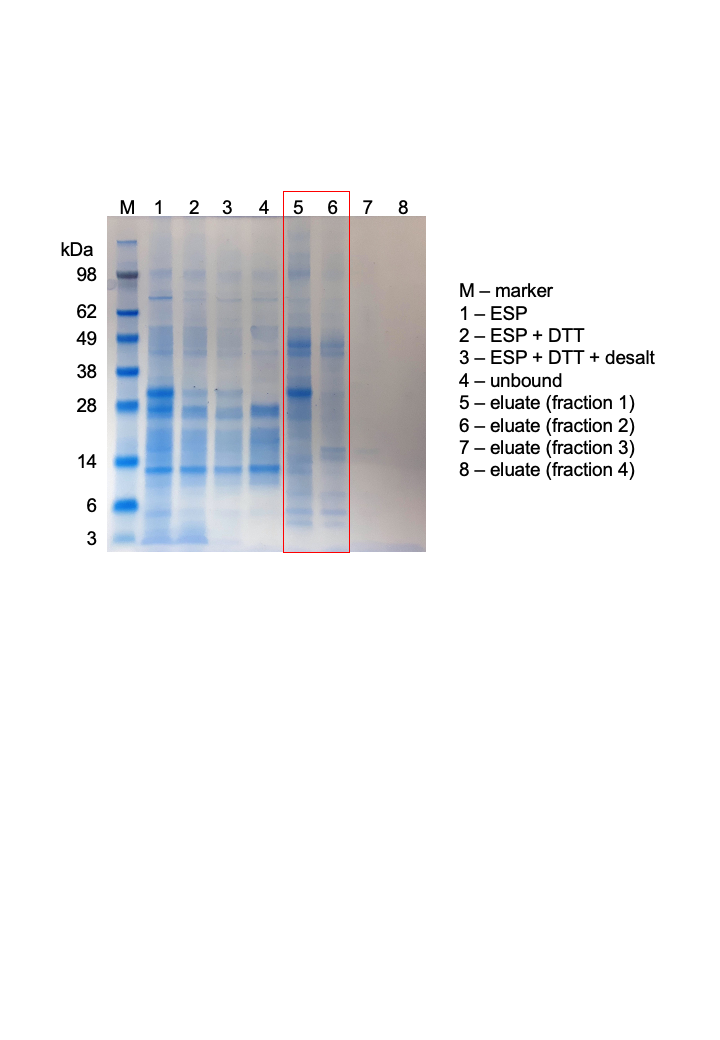

**Supplementary Figure 3.** SDS-PAGE analysis of *O. ostertagi* excretory-secretory proteins (ESPs) and purified ES-thiol under reducing conditions. Lanes 1-3 show unfractionated *O. ostertagi* adult ESPs. To purify ES-thiol, approx. 5mg of reduced and desalted *O. ostertagi* adult ESPs were loaded onto a thiol-sepharose column. Unbound material (lane 4) and elution fractions (lanes 5 – 8) are shown. Proteins contained in elution fractions 5 and 6 were pooled and used for proteomic analysis and vaccination-challenge studies in cattle.
