## Supplementary Table 1 for "Characterisation of protective vaccine antigens from the thiol-containing components of excretory/secretory material of *Ostertagia ostertagi*"

**Supplementary Table 1.** Summary of *O. ostertagi* ES-Thiol vaccine trials.

| **Trial** | **Breed** | **Sex** | **Age at V1 (days)** | **Group** | **n** | **Immunisation schedule** | **Challenge and post-mortem.** | **Post-mortem (day)** |
| --- | --- | --- | --- | --- | --- | --- | --- | --- |
| 1 | Holstein-Friesian | M/F | 156-166 | 1 | 6 | i.m. immunisation with 30 µg ES-Thiol + 750 µg Quil A^®^ (Brenntag Biosector) on days 0, 21 and 42. | Oral challenge with 1,000 *O. ostertagi* L3 per day, 5 days/week from days 42 to 76. | 91 |
|  |  |  |  | 2 | 6 | i.m. immunisation with 750 µg Quil A^®^ (Brenntag Biosector) on days 0, 21 and 42. |  |  |
| 2 | Friesian / Norwegian Red | M | 155-207 | 1 | 10 | i.m. immunisation with 30 µg ES-Thiol + 750 µg Quil A^®^ (Brenntag Biosector) on days 0, 21 and 42. | Oral challenge with 1,000 *O. ostertagi* L3 per day, 5 days/week from days 42 to 76. | 98 |
|  |  |  |  | 2 | 10 | i.m. immunisation with 750 µg Quil A^®^ (Brenntag Biosector) on days 0, 21 and 42. |  |  |
| 3 | Holstein/  Holstein-Friesian | M | 149-223 | 1 | 14 | s.c. immunisation with 15 µg ES-Thiol + 1 mg Vax Saponin^®^ (Guinness) on days 0, 21 and 42. | Oral challenge with 50,000 *O. ostertagi* L3 on day 42. | 76 |
|  |  |  |  | 2 | 14 | s.c. immunisation with 1 mg Vax Saponin^®^ (Guinness) on days 0, 21 and 42. |  |  |
| 4 | Mixed* | M/F | 114-161 | 1 | 12 | s.c. immunisation with 15 µg ES-Thiol + 1 mg Vax Saponin^®^ (Guinness) on days 0, 21 and 42. | Oral challenge with 50,000 *O. ostertagi* L3 on day 42. | 76 |
|  |  |  |  | 2 | 12 | s.c. immunisation with 1 mg Vax Saponin^®^ (Guinness) on days 0, 21 and 42. |  |  |

i.m. = intra-muscular; s.c. = subcutaneous; M = male; F = female

*Mix of Stabiliser crossbreed; British Blue crossbreed; Aberdeen Angus crossbreed; Holstein-Friesian; Limousin crossbreed; Hereford crossbreed; Norwegian Red crossbreed.
